# Co-delivery of human adipose-derived stromal cells and endothelial colony-forming cells in cell-assembled decellularized adipose tissue scaffolds for applications in soft tissue regeneration

**DOI:** 10.1101/2025.08.21.671493

**Authors:** Sarah A. From, John T. Walker, Connor J. Gillis, John A. Ronald, David A. Hess, Lauren E. Flynn

## Abstract

Cellular therapies involving the co-delivery of cells with complementary pro-regenerative functionality hold promise as a strategy to promote soft tissue regeneration. In particular, the co-delivery of adipose-derived stromal cells (ASCs) and endothelial colony-forming cells (ECFCs) has shown promise for regenerating stable blood vessels *in vivo.* The current study developed novel “cell-assembled” scaffolds for co-delivering human ASCs and ECFCs within a supportive decellularized adipose tissue (DAT) matrix, with the objective of enhancing their localized retention and augmenting their capacity to stimulate adipose tissue regeneration. Human ASCs and ECFCs were seeded separately onto human-derived DAT microcarriers under cell-type specific conditions. The cell-seeded microcarriers were then combined and cultured for 8 days under conditions that promoted matrix remodeling to fuse the microcarriers into 3D engineered tissues containing ASCs+ECFCs, ASCs alone, or ECFCs alone. Co-culture with ECFCs within the scaffolds was shown to modulate ASC pro-angiogenic gene expression, with some ECFCs forming tubule-like structures *in vitro* in both the ASC+ECFC and ECFC alone groups. *In vivo* bioluminescence imaging using a dual luciferase reporter system showed that co-delivery with ASCs enhanced ECFC retention following subcutaneous implantation in athymic *nu/nu* mice, but co-delivery did not alter the localized retention of viable ASCs. Interestingly, while immunofluorescence staining for CD31 and microcomputed tomography angiography indicated that vascular regeneration was similar in the cell-assembled scaffolds containing ASC+ECFCs, ASCs alone, and ECFCs alone, histological staining revealed that extensive regions of the ECFC alone scaffolds had remodelled into adipose tissue at 29 days post-implantation.

## 1 Introduction

Damage or loss of the subcutaneous adipose tissue layer as a result of oncologic resection, congenital defects, trauma, and burns can result in scar tissue formation, contracture, and deformities [1,2]. Patients with such soft tissue defects can often experience anxiety, depression, and physical disabilities, and successful reconstruction can profoundly improve their quality of life [3]. However, long-term stability and volume retention using techniques such as autologous tissue transfer are difficult to achieve clinically [4].

Decellularization of allogeneic tissues can be used to create “off-the-shelf” scaffolds that can stimulate the regeneration of damaged tissues and hold promise for a diverse range of clinical applications [5–7]. In the context of adipose tissue engineering, decellularized adipose tissue (DAT) has similar mechanical properties to native human fat [8] and is enriched in pro-regenerative extracellular matrix (ECM) components [9–11]. Numerous studies have reported that DAT processed into a variety of scaffold formats has pro-adipogenic effects both *in vitro* and *in vivo* [9,12,13]. In addition, we have demonstrated that human fibroblast survival and pro-angiogenic function were enhanced on DAT foams as compared to structurally-similar collagen foams [10], further supporting the use of DAT to develop pro-regenerative cell delivery platforms.

While applying DAT as an off-the-shelf scaffold is attractive, the rate and extent of adipose tissue regeneration can be enhanced by seeding the scaffolds with regenerative cells [14–16], which may be critical for larger soft tissue defects. Human adipose-derived stromal cells (ASCs) are abundant within fat, are highly tolerant of ischemic conditions [17], and show enhanced adipogenic potential and more potent immunomodulatory effects than bone marrow-derived mesenchymal stromal cells (MSCs) [18–20]. A limitation of traditional top-down tissue engineering approaches for scaffold fabrication for larger-volume applications is that they often result in cell localization primarily in the scaffold periphery unless technically-sophisticated bioreactor approaches are applied [21,22]. To address this limitation, we recently established a new bottom-up method of fabricating “cell-assembled” DAT scaffolds that contain a high density and even distribution of viable ASCs [23]. The scaffolds were generated by culturing ASC-seeded human-derived DAT microcarriers under conditions that promote cell expansion and ECM remodeling to fuse the microcarriers with new cell-secreted ECM into robust engineered tissues that maintain the geometry of the moulds used for fabrication. The cell-assembly process had a preconditioning effect on the human ASCs within the scaffolds, augmenting their survival and pro-angiogenic functionality [23].

While the delivery of ASCs alone may be beneficial, recent evidence supports that combined cell therapies may help to accelerate and augment regeneration [24–27]. Complementing the pro-regenerative functionality of ASCs, endothelial colony-forming cells (ECFCs) are endothelial precursor cells that can be derived from umbilical cord blood or adult bone marrow [28], and have the capacity to form *de novo* vessels *in vivo* [28–30]. ECFCs have also been reported to secrete a range of paracrine factors involved in blood vessel formation [24,31,32]. Studies support that ASCs and ECFCs can communicate through direct cell-cell interactions, as well as via paracrine mechanisms, and that they may have complementary pro-angiogenic secretomes [24,33,34]. ASCs can also function as vessel-stabilizing cells to support new blood vessels formed by ECFCs in areas of ischemia [34]. Similarly, the ECFC secretome has been reported to enhance ASC engraftment and adipogenic differentiation *in vivo*, as well as helping to counter their senescence [24].

Recognizing the potential of combination cell therapies, our objective in this study was to adapt our cell-assembly methods to generate scaffolds incorporating both human ASCs and ECFCs and characterize their properties *in vitro* relative to cell-assembled scaffolds fabricated with ASCs alone or ECFCs alone. To more fully assess the potential benefits of our new combined cell therapy platform, *in vivo* studies were performed using a subcutaneous implantation model in athymic *nu/nu* mice [35] to assess the effects of co-delivery on the localized retention of viable ASCs and/or ECFCs, as well as on markers of angiogenesis and soft tissue regeneration.

## 2 Methods

### 2.1 Human adipose tissue collection and ASC isolation and culture

Adipose tissue samples were collected with informed consent from elective lipo-reduction and breast reconstruction surgeries with Research Ethics Board approval from Western University (HSREB # 105426). All adipose tissue samples were processed within 2 h of isolation using published protocols for either decellularization [44] or ASC isolation [45]. ASCs were cultured in Dulbecco’s modified Eagle’s medium/Ham’s F12 (DMEM/F12) supplemented with 10% fetal bovine serum (FBS), 100 U/mL penicillin, and 0.1 mg/mL streptomycin at 37°C and 5% CO_2_, with media changes every 2-3 days and passaging at ∼80% confluence. ASCs at passage 3-4 were used for scaffold fabrication.

### 2.2 Human endothelial colony-forming cell (ECFC) isolation and culture

Human umbilical cord blood samples were collected with informed consent from scheduled cesarian section births (HSREB # 12934) in 0.5 mL heparin and processed within 24 h following published methods [36]. ECFCs were cultured in endothelial growth medium (EGM-2 + Bullet Kit, Lonza), and passaged at ∼80% confluence. ECFCs at passage 4-6 were used for scaffold fabrication.

### 2.3 Cell transductions

To enable cell tracking of both cell populations, published lentiviral transduction methods were adapted [37] to engineer the ASCs to express the reporters tdTomato (tdT) and firefly luciferase (Fluc2). Similarly, ECFCs were transduced to express miRFP720 and the bioluminescence resonance energy transfer (BRET)-based Antares luciferase reporter [38]. Cells were transduced at a multiplicity of infection (MOI) of 2, based on functional titre, with 8 μg/mL polybrene added to the culture media. Fluorescence-activated cell sorting (FACS) was performed (FACSAria™ III, BD Biosciences) to enrich the transduced ASC population. Transduced cells were used for all *in vivo* work, as well as for the *in vitro* flow cytometry, histology, and RT-qPCR studies.

### 2.4 DAT microcarrier fabrication

Published electrospraying methods were applied to generate DAT microcarriers that formed the subunits of the cell-assembled scaffolds [39]. In brief, DAT (pooled from a minimum of 5 donors) was lyophilized and cryomilled using a Retsch MM 400 mixer mill prior to enzymatic digestion with 1% w/w α-amylase in 0.22 M NaH_2_PO_4_ in deionized water (diH_2_O) on an orbital shaker at 300 rpm for 72 h at room temperature (RT). The DAT was homogenized in 0.2 M acetic acid to generate a 35 mg/mL ECM suspension that was incubated at 37°C under agitation at 100 rpm overnight. The DAT suspension was electrosprayed at a rate of 30 mL/h into liquid nitrogen at 10-20 kV through a 25 G winged infusion set (BD Medical, Mississauga, ON, Canada), with the needle tip positioned 5 cm from the surface of the liquid nitrogen. The microcarriers were collected and lyophilized for ∼24 h. To prevent loss of structure due to swelling, the microcarriers were gradually rehydrated through an ethanol in PBS series (99, 98, 95, 90, 80, 70, 60, 50, 40, 30, 20, 10, 0% ethanol in PBS; 30 min/step, incubation under agitation at 200 rpm and RT) and then stored in 100% sterile PBS at 4°C.

### 2.5 Dynamic cell seeding on DAT microcarriers

An overview of the cell-assembly protocol is shown in Figure 1A. Published methods [40] using 100 mL CELLSPIN spinner flasks were adapted to seed the ASCs and ECFCs on the DAT microcarriers in their respective culture media described above. The ASCs were seeded at 1.25×10^6^ ASCs/g of microcarriers (wet weight in 100% EtOH) and the ECFCs were seeded at 2.5 x10^6^ ECFCs/g of microcarriers, based on preliminary testing that indicated a higher seeding density was required for the ECFCs to achieve a homogenous distribution across the microcarrier surface (Supplementary Figure S1). The media was topped up to 100 mL and the flasks were subjected to a 24 h stirring regimen at 37°C and 5% CO_2_ as follows: (i) 2 min at 25 rpm followed by 30 min at 0 rpm, repeated 6 times, followed by (ii) 6 h at 0 rpm, an additional cycle of (iii) 2 min at 25 rpm followed by 30 min at 0 rpm, repeated 6 times, and finally, (iv) 12 h of continuous stirring at 25 rpm.

**Figure 1.**
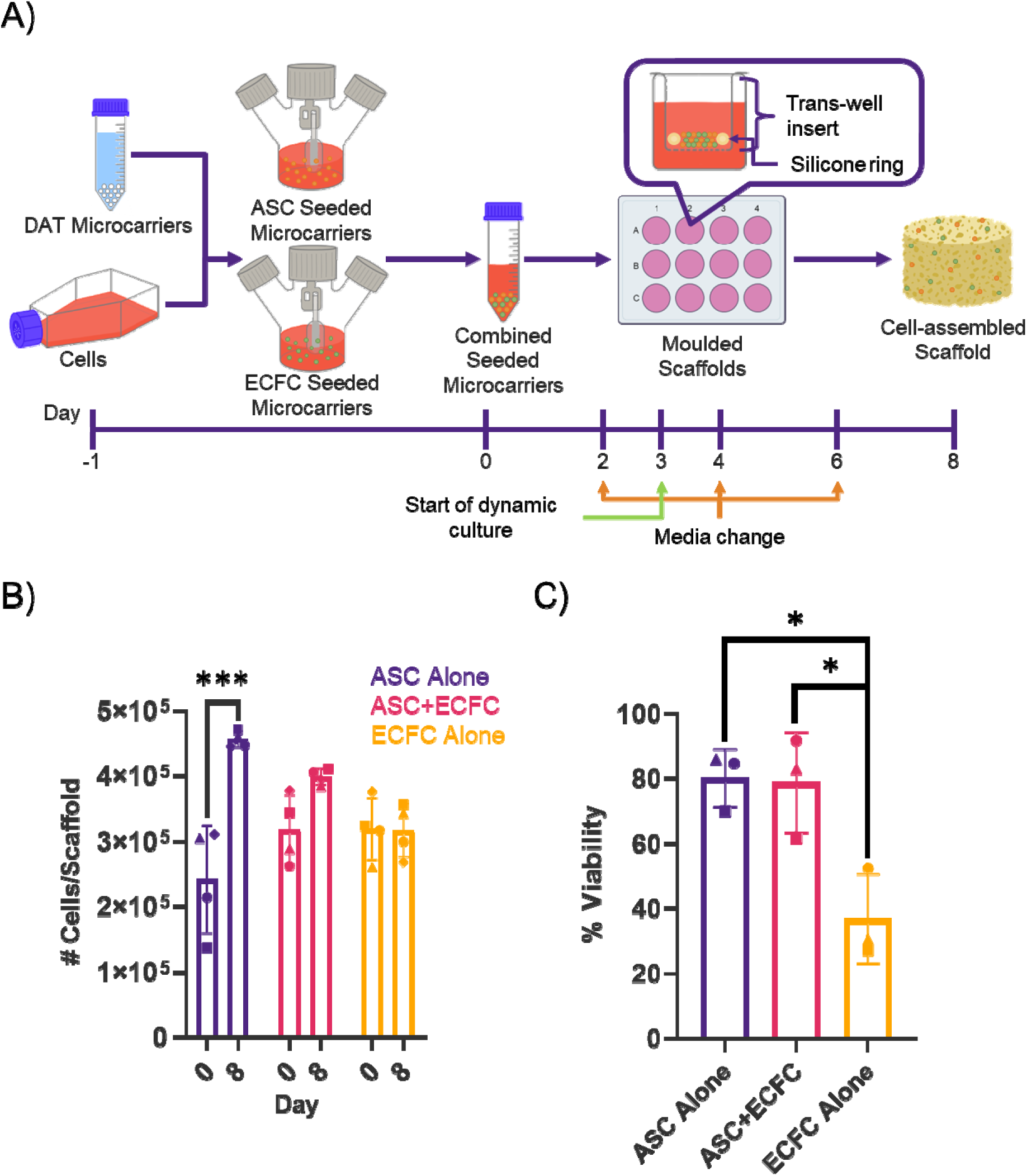
Cell-assembled scaffolds were generated with human ASCs+ECFCs, ASCs alone and ECFCs alone that contained similar numbers of cells, but cell viability was lower in the ECFC alone group. (A) Schematic overview of the cell-assembly process, involving the seeding of DAT microcarriers with ASCs and ECFCs in separate spinner flask bioreactors over 24 h, followed by the transfer of the seeded microcarriers into moulds, where they were cultured for an additional 8 days to generate the cell-assembled scaffolds. (B) Quantification of cell abundance via the PicoGreen assay at days 0 and 8 (n=3, N=4). (C) Flow cytometric analysis of cell viability at day 8 (n=3, N=3). *p<0.05, ***p<0.005.

### 2.5 Live/Dead® imaging

Cell viability and attachment to the microcarriers were visually assessed after the 24 h spinner flask seeding regimen using a Live/Dead® assay kit (Life Technologies) [41] and a Zeiss LSM confocal microscope (Germany). Z-stacks (10 images/sample) through the microcarriers were created at 5 X magnification.

### 2.6 Cell-assembled scaffold fabrication and culture

Adapting published methods [23], the cell-seeded microcarriers were transferred into moulds constructed using 12-well plates with ThinCert® cell culture inserts (Greiner Bio-one) lined with medical grade silicone rings (8 mm inner diameter, 12 mm outer diameter, 3 mm height). Scaffolds were generated with (i) ASC-seeded microcarriers alone, (ii) ECFC-seeded microcarriers alone, or (iii) a 1:1 ratio of ASC:ECFC-seeded microcarriers. All samples were cultured (37°C, 5% CO_2_) in the moulds in Prime-XV® MSC EXPANSION XSFM Medium (Irvine Scientific) supplemented with the EGM-2 BulletKit™ (Lonza), based on preliminary testing that showed this formulation supported the viability of both cell populations, with media changes every 2 days. After 3 days of static culture, the samples were cultured dynamically on an orbital shaker at 75 RPM for 5 additional days.

### 2.7 PicoGreen™ quantification of dsDNA content

To quantify total cell abundance within the scaffolds at day 0 (post-moulding) and at the end of the 8-day assembly phase, samples were processed with the DNeasy Blood & Tissue Kit (Qiagen) prior to analysis with the Quant-iT™ PicoGreen™ assay (Thermo Fischer Scientific) using a CLARIOstar® microplate reader [42,43]. The dsDNA content was converted to cell number by normalization to standard curves generated with known numbers of ASCs and/or ECFCs.

### 2.8 Flow cytometric analysis of cell viability

Flow cytometry analyses were performed to assess cell viability in the scaffolds at the end of the 8-day assembly phase. The scaffolds were minced and enzymatically digested with 1000 U/mL collagenase type IV in a 1:1 mixture of EBM-2 and DMEM/F12 supplemented with 100 U/mL penicillin, 0.1 mg/mL streptomycin for 2 h at 37°C and 75 rpm. After enzymatic digestion, the collagenase was inactivated with an equal volume of ASC growth medium and the samples were filtered through a 70 µm cell strainer, centrifuged at 350 x g for 7 min, and resuspended in PBS+5% FBS. The cells were stained with 10 nM CellTracker™ Green CMFDA (Thermo Fisher) in DMEM/F12 for 45 min at 37°C, and washed for 30 min in PBS+5% FBS at 37°C prior to analysis on an LSR II Flow Cytometer (BD Biosciences). Gating was performed using fluorescence minus one (FMO) controls. Singlets were identified based on forward versus side scatter profiles and live cells were identified using CellTracker™ Green. Data analyses were performed using FlowJo software.

### 2.9 Analysis of cell distribution

Samples were cryo-sectioned to analyze the distribution of the transduced ASCs and ECFCs within the scaffolds based on the expression of the fluorescent reporter genes. Samples were fixed through overnight incubation in 10% neutral buffered formalin. Next, the samples were incubated overnight in 15% sucrose and then 30% sucrose solutions at 4°C, and then embedded in VWR® clear frozen section compound for cryosectioning (12 µm sections). Sections were mounted in DAPI-enhanced mounting medium (abcam) and imaged using an EVOS FL fluorescence microscope (Thermo Fisher Scientific Inc.) at 10 X magnification. A minimum of 6 non-overlapping fields of view were imaged per slide, with 3 slides taken from different depths within each scaffold.

### 2.10 Biochemical analysis of matrix composition

The hydroxyproline and dimethylmethylene blue (DMMB) assays were performed to quantitatively compare the total collagen and sulphated glycosaminoglycan (GAG) content between the ASC alone, ECFC alone and ASC+ECFC scaffolds at days 0 and 8, following published protocols [44]. For the hydroxyproline assay, the absorbance was measured at 560 nm and for the DMMB assay, the absorbance was measured at 525 nm and corrected at 595 nm, using a CLARIOstar® microplate reader.

### 2.11 Angiogenic factor PCR array

The Human Angiogenic Growth Factor RT² Profiler™ PCR Array (Qiagen) was used to compare the transcriptional profile of ASCs isolated from the ASC+ECFC scaffolds to the ASC alone scaffolds at the end of the 8-day assembly phase. In both groups, the ASCs were isolated using FACS based on tdT fluorescence expression, as described above. Total RNA was extracted using PureZOL™ and the samples were then processed with the Aurum™ Total RNA Fatty and Fibrous Tissue Kit (Bio-Rad) following the manufacturer’s instructions. For each sample, 400 ng of RNA was processed into cDNA using the Qiagen RT^2^ First Strand Kit. The PCR Array was performed following the manufacturer’s instructions using RT^2^ SYBR® Green qPCR Mastermix. Data was normalized to the geometric mean of 5 housekeeping genes (beta actin; beta-2-microglobulin; glyceraldehyde-3-phosphate dehydrogenase; hypoxanthine phosphoribosyl transferase 1; Ribosomal protein, large, P0), analyzed using the 2^-ΔCt^ method, and reported as Log_2_ fold change.

### 2.12 Scaffold implantation in athymic nu/nu mice

*In vivo* studies were performed to compare cell retention and scaffold remodeling in the combined ASC+ECFC scaffolds to ASC alone and ECFC alone controls using a subcutaneous implant model in 8-10 week-old athymic nude mice (Crl:NU-Foxn1nu; Strain Code 088, Charles River, Wilmington, MA, USA). All studies complied with the Canadian Council on Animal Care guidelines and were approved by the Animal Care Committee at Western University (AUP #2023-102). Using published protocols [35], two scaffolds of the same type were implanted subcutaneously in separate pockets on the dorsa of each mouse. A total of 12 mice (6 male, 6 female) were used for bioluminescence imaging (BLI) and an additional 16 mice (8 Female, 8 male) were used for microcomputed tomography (μCT) angiography, described below.

### 2.13 In vivo analysis of cell retention and host endothelial cell recruitment

Longitudinal *in vivo* cell tracking was performed using BLI to quantify viable FLuc^+^ ASCs and Antares^+^ ECFCs over 29 days post-implantation. Imaging of the two populations was performed 24 h apart to ensure no overlap in the signals. On days 1, 3, 7, 14, 21, and 28, mice were anesthetized using isoflurane and received an intraperitoneal (IP) injection of 30 mg/mL D-luciferin in PBS (100 µL) for ASC tracking. Similarly, on days 2, 4, 8, 15, 22, and 29, mice were treated with a 50 µL IP injection of 3.1 μM Nano-Glo® fluorofurimazine (FFz) in PBS (Promega) for ECFC tracking. BLI was performed using an IVIS Lumina XRMS *In Vivo* Imaging System, as previously described [23]. Mice were imaged once every minute, and a consistently sized region of interest (ROI) was drawn around the luminescent signal to quantify the average radiance (p/s/cm^2^/sr) until the signal peaked. Peak values were normalized to the day 1 or 2 values, for ASCs and ECFCs respectively, for each mouse.

Mice were euthanized following the day 29 imaging timepoint, and the scaffolds were excised within their surrounding tissues and cryo-sectioned. Immunofluorescence staining was performed to identify CD31^+^ endothelial cells within the implants. Sections were blocked in PBS+10% donkey serum for 2 h at RT before the application of polyclonal goat anti-CD31 primary antibody (R&D Systems, AF3628) at a 1:40 dilution in PBS+10% donkey serum and incubation overnight at 4°C. The slides were then rinsed in PBS and donkey anti-goat IgG secondary antibody (Invitrogen, A11055) was applied at a 1:500 dilution in PBS+10% donkey serum and incubated for 2 h at RT. The slides were rinsed with PBS and mounted in DAPI mounting medium prior to imaging on an EVOS FL fluorescence microscope, with 6 random, non-overlapping fields of view captured for each cross-section. Positive pixel counting was performed within the scaffold region using Fiji software, and the CD31 signal was normalized to the scaffold area. High-resolution images were also taken on a Zeiss LSM confocal microscope at 25 X magnification.

### 2.14 Microcomputed tomography angiography

A separate group of mice (N=16, 8 female, 8 male) was used for μCT angiography to assess the perfused vasculature within and surrounding the implants after 28 days *in vivo*. Unseeded scaffolds [35] were included as an imaging control group to validate our previous findings that showed vessel leakage in this group [23]. Briefly, the mice were perfused using a gravity-fed perfusion system with MICROFIL® (Flow Teck Inc) contrast agent via a cannula inserted in the right atrium. The mice were fixed in 10% neutral buffered formalin and scans were acquired on a GE eXplore speCZT scanner (GE Healthcare, London, ON) as previously described [23].

Analysis was performed using MicroView (Parallax Innovations, London, ON, Canada) to locate the scaffolds and 3-D Slicer (http;//www.slicer.org) to measure the volume of the vessels and segment the scaffolds.

Following imaging, the implants were collected and further processed for histological analysis to assess tissue remodeling and the distribution of perfused blood vessels within the scaffolds.

Briefly, the scaffolds were embedded within their surrounding tissues in paraffin and sectioned (5 µm thickness). Sections were stained with hematoxylin and eosin (H&E) and scanned at 10× magnification using a Nikon Eclipse Ti2 microscope. Qualitative assessments were performed in a blinded manner to examine vessel size, infiltration, and the presence of adipocytes within the scaffolds.

### 2.15 Statistical analyses

All *in vitro* studies were repeated with ASCs and ECFCs from 3-4 different donors (N=3-4) and included a minimum of 2 to 3 replicate scaffolds per trial (n=2-3). For the *in vivo* studies, 2 cell donors were used with a minimum of 4 replicate scaffolds per scaffold group. Statistical analyses were performed using GraphPad Prism 8.4.3. All *in vitro* data is expressed as the mean ± standard deviation (SD) and analyzed by one-way or two-way ANOVA with Tukey’s or Sidak’s post-hoc comparison, unless otherwise indicated. For the *in vivo* studies, comparisons between groups were performed using a mixed effects model with Tukey’s or Sidak’s post-hoc comparison. Differences were considered statistically significant at p<0.05.

## 3 Results

### 3.1 Cell-assembled scaffolds were successfully generated containing both ASCs and ECFCs

Expanding on our cell-assembled platform, we successfully adapted our cell-assembly methods using multiple cell types for the first time. In particular, we generated robust scaffolds incorporating both human ASCs and ECFCs, as well as human ECFCs alone, in addition to our established ASC alone scaffolds. Capitalizing on our modular fabrication approach, the ASCs and ECFCs were seeded onto the DAT microcarriers under cell type-specific media conditions within separate spinner flask bioreactors (Figure 1A). Following 24 h seeding, the microcarriers were transferred, either alone or in a 1:1 ratio, into moulds and cultured for an additional 8 days to generate the ASC alone, ECFC alone and ASC+ECFC scaffolds.

The PicoGreen® assay was used to estimate the total cell abundance in the scaffolds at days 0 and 8 (Figure 1B). Consistent with our previous work [23], there was a significant increase in scaffold cellularity over the 8-day assembly phase in the scaffolds generated with human ASCs alone. In contrast, there was no significant change in the total cell content between days 0 and 8 for the scaffolds generated with ASCs+ECFCs or ECFCs alone. Regardless, the total cell abundance within the scaffolds was similar between all three groups at both day 0 and day 8.

Flow cytometry analysis of cell viability was performed on single cell suspensions prepared from collagenase-digested scaffolds at the end of the 8-day assembly phase (Figure 1C). Notably, cell viability was significantly reduced in the ECFC alone scaffolds (37 ± 14%) compared to the ASC alone (80 ± 9%) and ASC+ECFC scaffolds (79 ± 15%).

Cell distribution in the cell-assembled scaffolds at day 8 was visualized using scaffolds generated with ASCs and ECFCs engineered through lentiviral transduction to express the fluorescent reporters tdT and miRFP720, respectively (Figure 2). Cells were distributed throughout the scaffolds, with the ASCs showing a qualitatively higher density near the periphery of the scaffolds in both the ASC+ECFC and ASC alone groups. Notably, the ECFCs were observed to form tubule-like structures in both the ASC+ECFC and ECFC alone groups.

**Figure 2.**
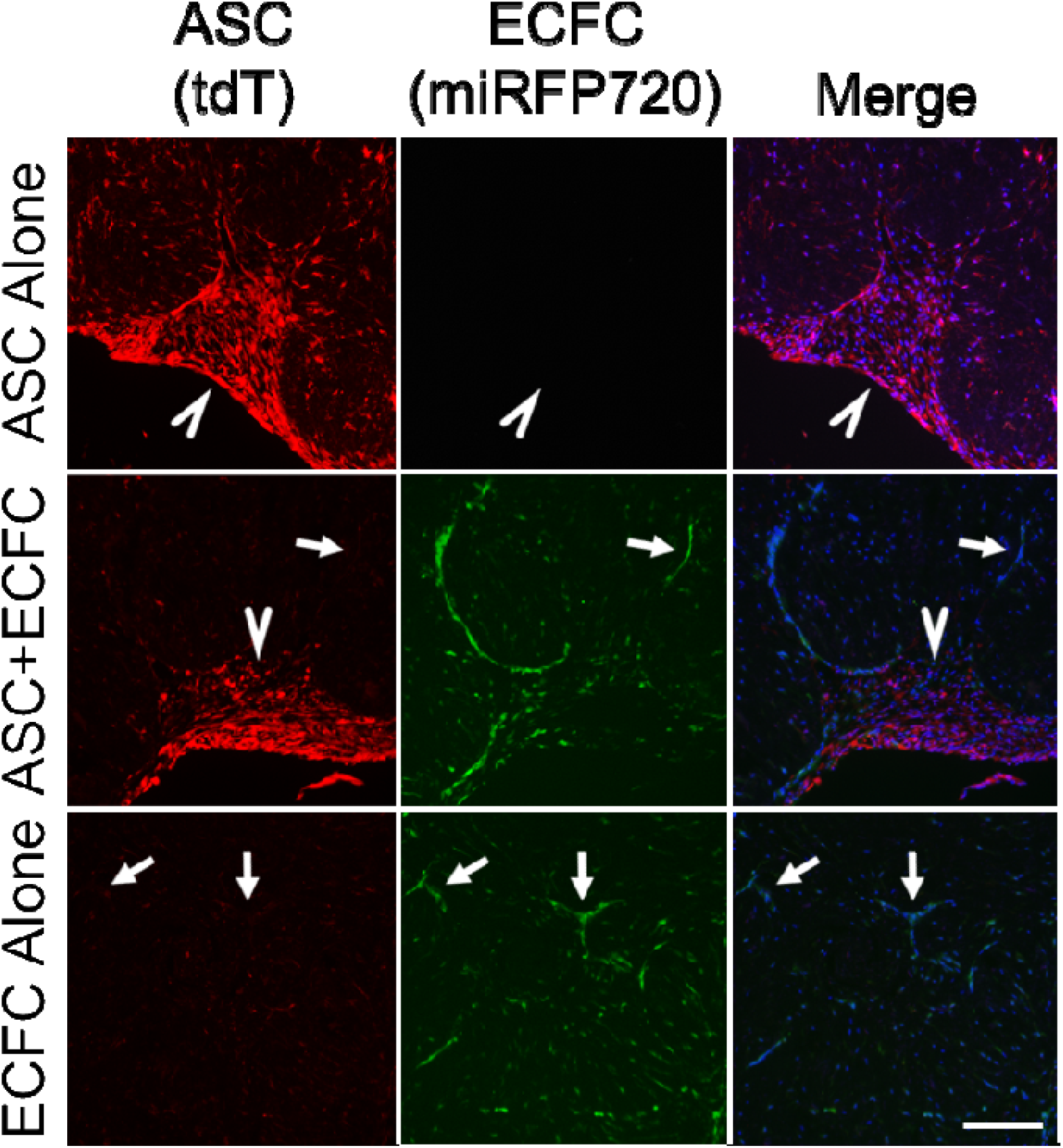
Cells were distributed throughout the cell-assembled scaffolds, with the ASCs having a higher density in the scaffold periphery and the ECFCs forming tubule-like structures in the scaffold interior. Representative images of cell-assembled scaffolds after the 8-day assembly phase. Arrow heads mark areas of high ASC density near the periphery of the scaffolds. Arrows mark tubule-like structures formed by ECFCs. DAPI staining is shown in blue Scale bar = 200 µm.

### 3.2 The cell assembly process was associated with matrix remodeling

Scaffold contraction was observed over the 8-day assembly phase, with no significant differences observed between the groups at day 8 based on surface area analysis (Figure 3A). However, there was greater donor variability in contraction in the ECFC alone group, with one donor showing markedly higher levels of contraction. Qualitatively, ECFC alone scaffolds that showed lower levels of contraction were softer and more fragile when being handled. However, compression testing confirmed that all scaffolds were soft and compliant, with no significant differences in the Young’s moduli between the groups at day 8 (Figure 3B). Notably, the Young’s modulus was highest for the ECFC alone scaffolds that were generated with the cell donor that showed higher levels of contraction (denoted by triangle symbol on plots).

**Figure 3.**
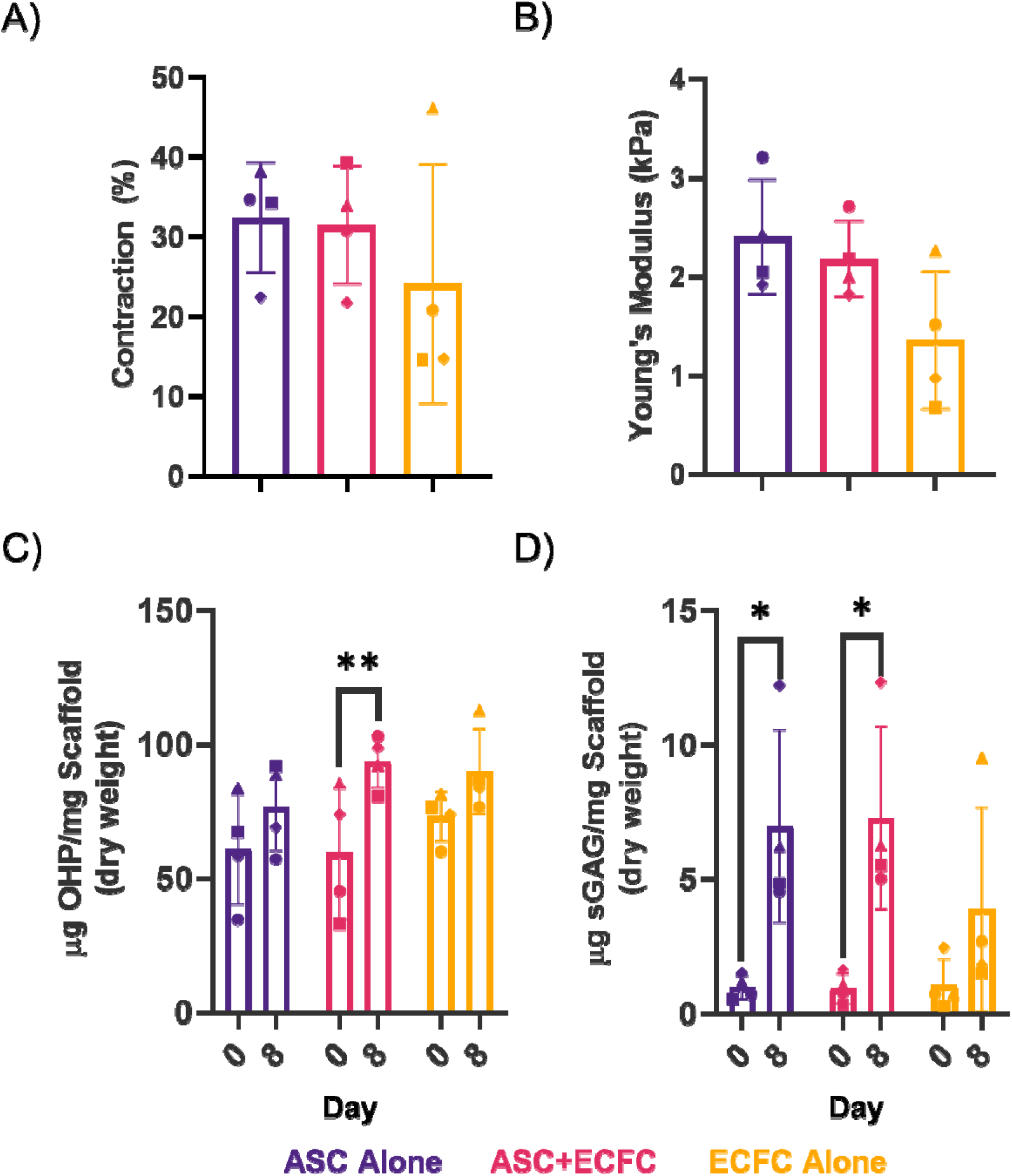
Cell-assembled scaffolds generated with ASCs and/or ECFCs were soft and compliant, with similar collagen and sulphated glycosaminoglycan content at the end of the 8-day assembly phase. (A) Scaffold contraction at day 8 based on analysis of scaffold surface area relative to the area of the mould (n=3, N=4). (B) Young’s modulus of the cell-assembled scaffolds measured at day 8 with a CellScale Univert compression testing system (n=2-3, N=4). C) Hydroxyproline content as a measure of total collagen content in the scaffolds at days 0 and 8 (n=3, N=4). D) Sulphated GAG content at days 0 and 8 measured using the DMMB assay. *p<0.05, **p<0.01.

Analysis of total collagen content via the hydroxyproline assay showed no differences between the scaffold groups at day 8, with the ASC+ECFC scaffolds showing a significant increase in hydroxyproline content from day 0 to 8, indicative of ECM remodeling (Figure 3C). Similarly, analysis of sulphated (GAG) content via the DMMB assay showed a significant increase in both scaffold groups containing ASCs over time (Figure 3D).

### 3.3 The presence of ECFCs within the cell-assembled scaffolds altered ASC angiogenic gene expression

An RT-qPCR array was used to compare the gene expression levels of 84 different angiogenesis-related genes in FACS-isolated tdT^+^ ASCs extracted from the ASC alone scaffolds and the combined ASC+ECFC scaffolds at the end of the 8-day assembly phase. Figure 4 presents the Log_2_ fold change of all genes that were consistently upregulated or downregulated in the scaffolds generated with three different cell donors. Based on multiple t-tests with a false discovery rate of 10%, ANGPT2 and RUNX1 were significantly upregulated (*p<0.05) and TIMP1 and CXCL13 were significantly downregulated in the ASCs from the combined ASC+ECFC scaffolds compared to the ASC alone scaffolds. Taken together, these findings suggest that co-assembly with the ECFCs modulated the pro-angiogenic phenotype of the ASCs.

**Figure 4.**
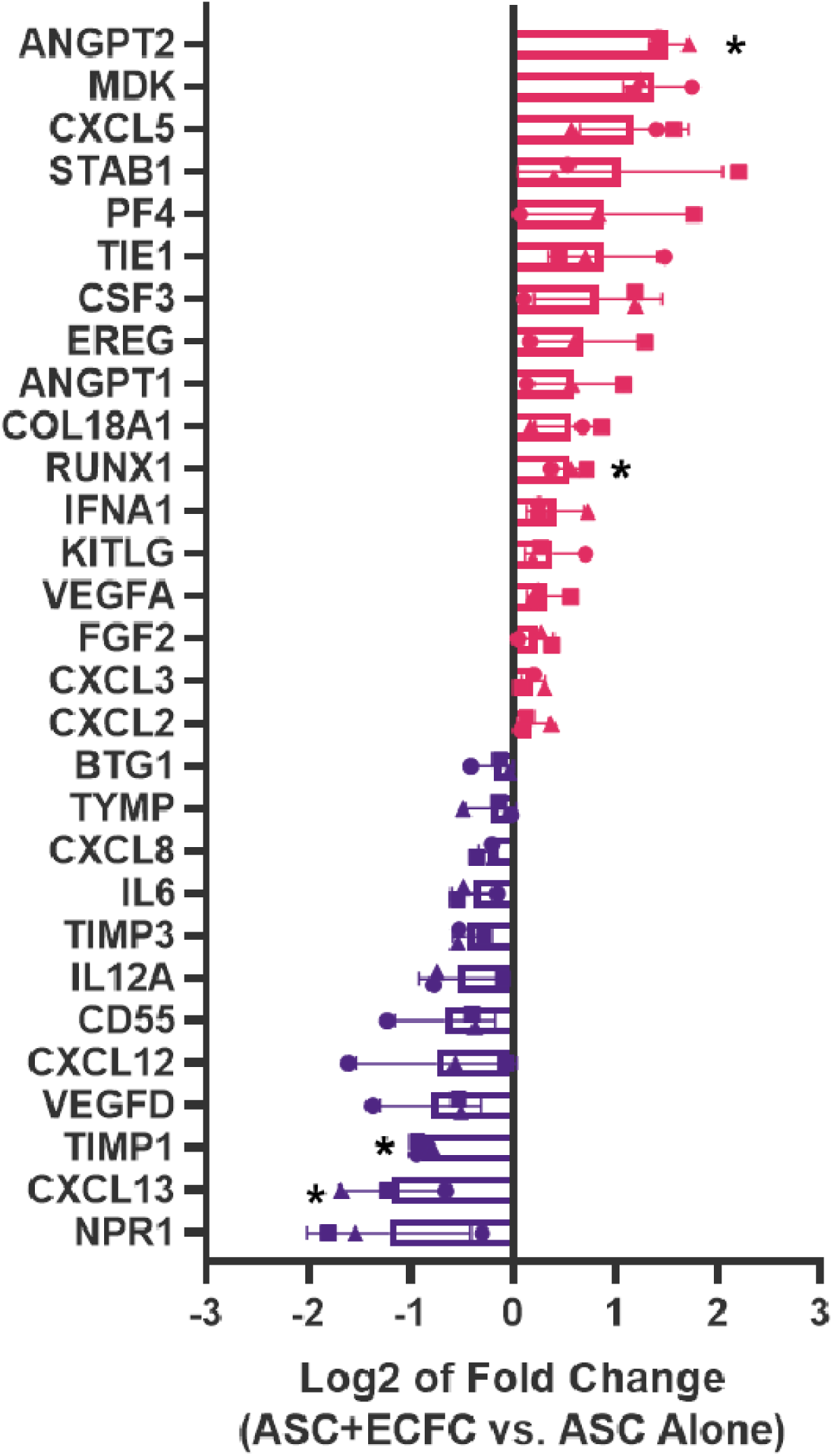
Co-culture with ECFCs in the cell-assembled scaffolds modulated angiogenesis-related gene expression in the ASCs. ASC gene expression was compared in cells isolated from the ASC+ECFC versus ASC alone scaffolds using an angiogenic factor qPCR array. Data presented as Log_2_ fold-change of mRNA levels of genes consistently up- or down-regulated across all 3 cell donors. (n=3 replicate scaffolds/donor (pooled), N=3; multiple t-tests with a false discovery rate of 10%).

### 3.4 Co-delivery with ASCs improved viable ECFC retention within the scaffolds following subcutaneous implantation in athymic nude mice

Following *in vitro* characterization, *in vivo* studies were performed to compare cell retention and tissue remodeling in the cell-assembled ASC+ECFC scaffolds to ASC alone and ECFC alone controls following subcutaneous implantation in athymic nude (*nu/nu*) mice. The ASCs and ECFCs were engineered to express the FLuc and Antares reporters respectively, to enable longitudinal tracking of both cell populations using orthogonal BLI substrates. The luminescent signal from the ASCs and ECFCs remained localized in the flank and visible over the 29-day testing period (Figure 5A,B). Based on quantitative comparison of the ASC signal over time (normalized to day 1), there was no significant difference between the ASC+ECFC scaffolds compared to the ASC alone controls, indicating that the localized retention of viable ASCs was similar in both groups (Figure 5C). In contrast, a significantly higher ECFC signal was detected in the combined ASC+ECFC scaffolds compared to the ECFC alone scaffolds at day 29, suggesting that the presence of the ASCs enhanced the retention of viable ECFCs over time (Figure 5D).

**Figure 5.**
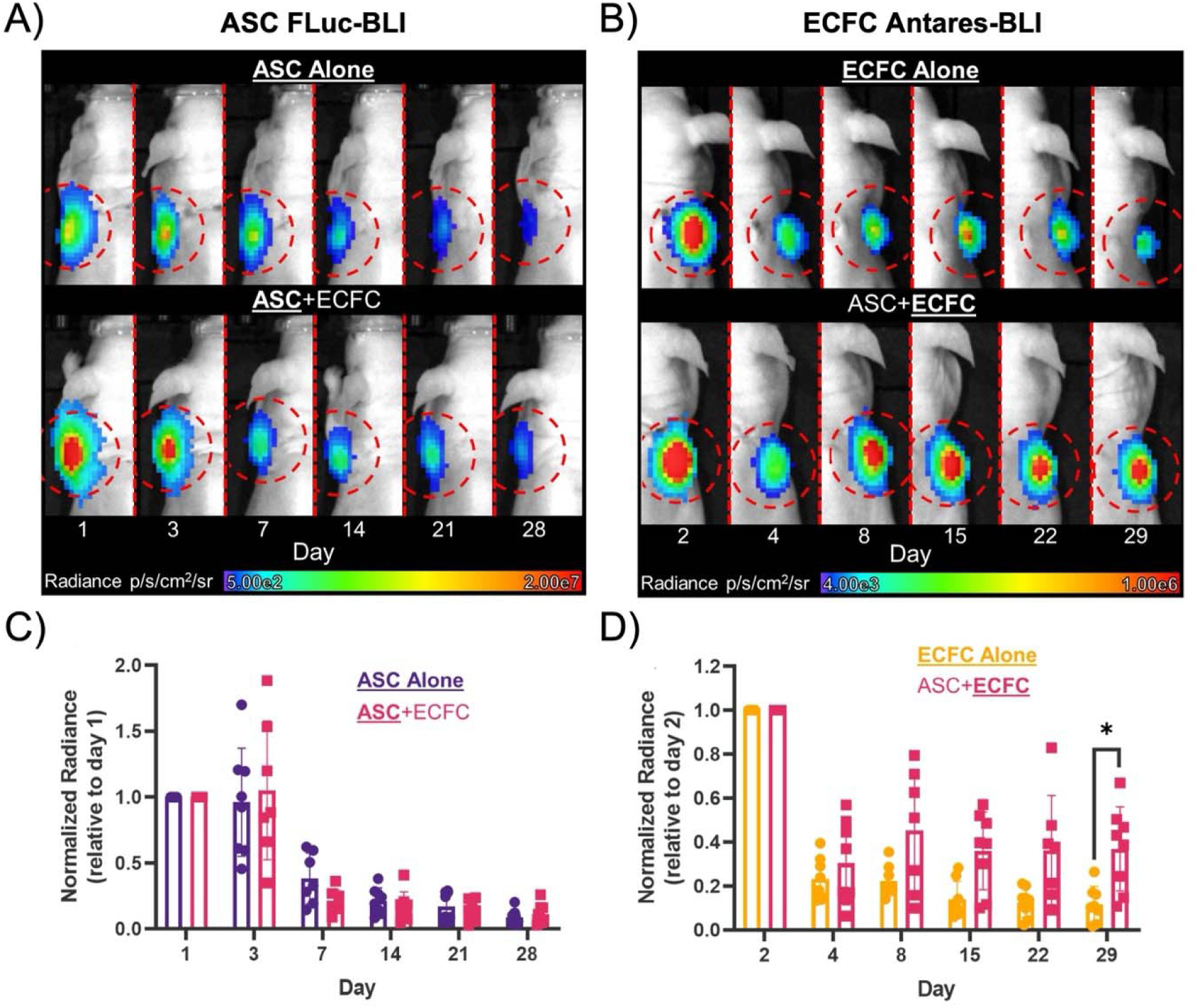
Co-delivery with ASCs enhanced the localized retention of viable ECFCs within the cell-assembled scaffolds *in vivo*. Representative images showing the peak BLI signal associated with the (A) ASCs or (B) ECFCs within the scaffolds. Red circles represent the measured ROI. Relative radiance associated with viable (C) ASCs normalized to day 1 and (D) ECFCs normalized to day 2. Statistical analyses were performed using a mixed effects model (N=7-8). *p<0.05.

### 3.5 CD31^+^ vessel density was similar between implanted scaffold groups

At day 29, the mice were euthanized following BLI and the tissues in the implant region were collected and processed for immunofluorescence staining to assess CD31^+^ host cell infiltration into the scaffolds. In general, the scaffolds were well integrated with their surrounding tissues, with elongated vessel-like CD31^+^ structures and lumens observed within the scaffold boundaries in all groups (Figure 6). Positive pixel counting showed no difference in the CD31 positive pixels per scaffold area between the scaffold groups (Supplementary Figure S2).

**Figure 6.**
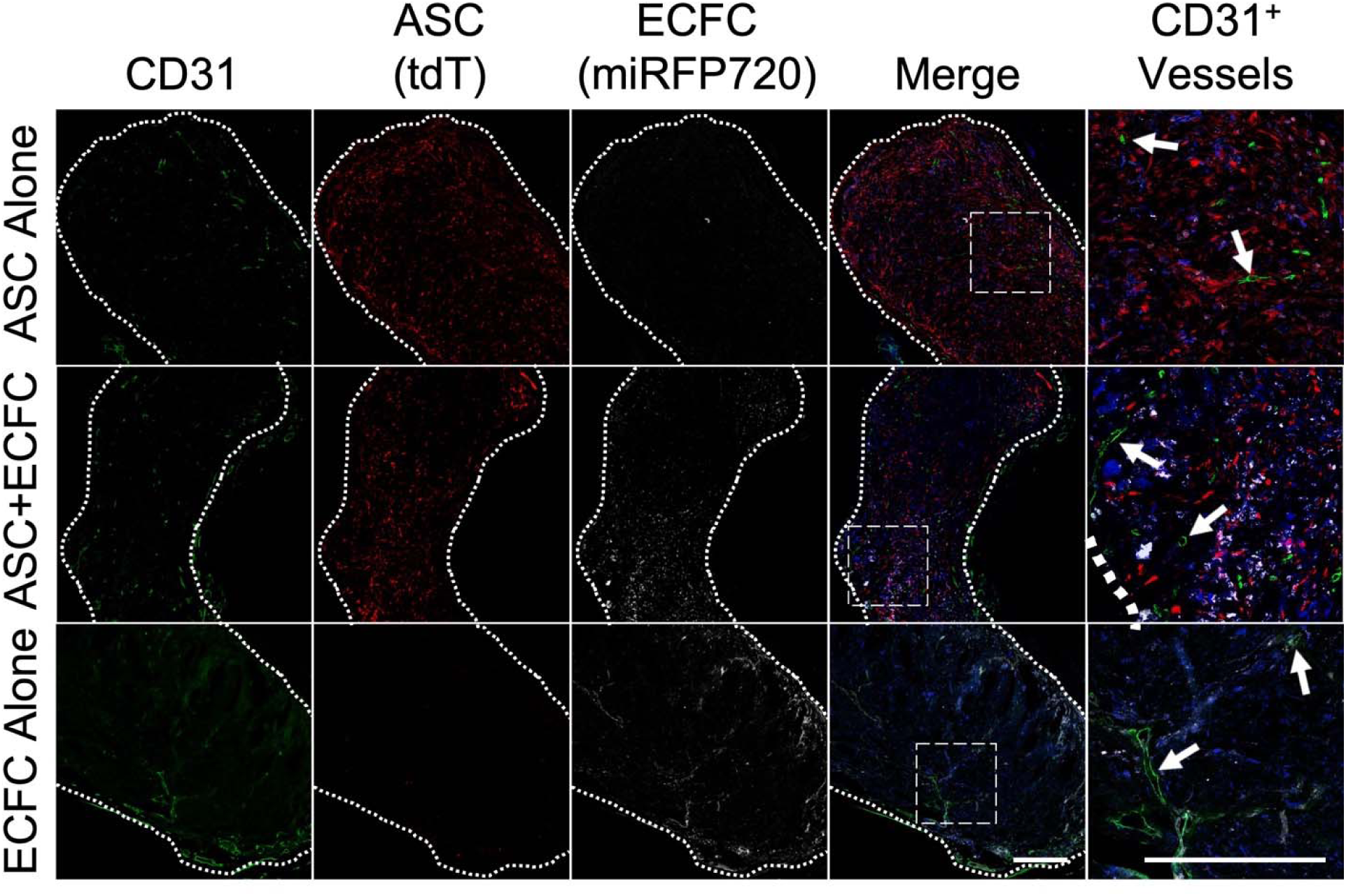
CD31^+^ host cells infiltrated into the scaffolds after 29 days *in vivo*. Representative images of scaffold cross-sections, showing the separate channels to localize the transplanted ASCs and/or ECFCs, with dotted lines indicating the scaffold perimeter. Higher magnification images of the boxed regions are shown on the right. Arrows indicate CD31^+^ lumens, observed in all scaffold groups. DAPI staining is shown in blue. Scale bars = 300 µm.

### 3.6 MicroCT angiography showed that the perfused vessel volume in the scaffold region was similar between the groups at day 29

At day 29, μCT angiography was performed on a separate cohort of mice to assess whether there were differences in the perfused vascular network structure in the implant region. Perfused vessels were observed both at the surface and infiltrating the scaffolds in all groups (Figure 7A). Quantification of the perfused vessel volume, as well as the total scaffold volume, showed no significant differences between the groups (Figure 7B,C). While there was leaking of the contrast agent observed in the unseeded scaffold controls (Supplementary Figure S3), no leaking was observed in any of the cell-assembled scaffold groups, suggesting that the incorporation of both cell populations was beneficial for reducing vascular permeability by accelerating angiogenesis and/or promoting vessel maturation.

**Figure 7.**
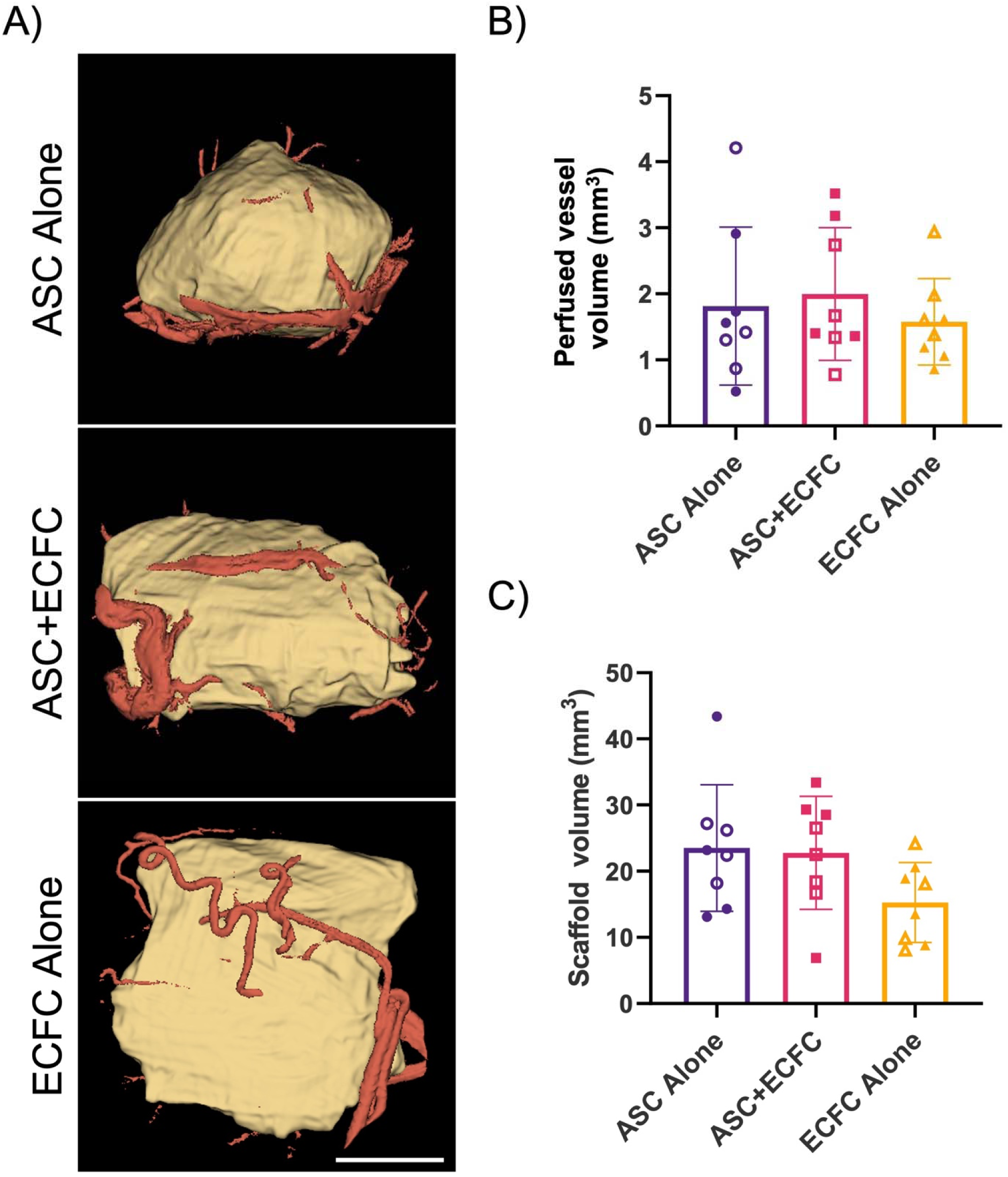
Similar levels of perfused blood vessels were observed in proximity to all cell-assembled scaffold groups at day 29. (A) Representative images of perfused vessels (red) in proximity to the scaffolds (beige) after 29 days *in vivo*. Scale bar = 2.5 mm. (B) Average volume of perfused vessels within 0.5 mm of scaffold surface. (C) Average scaffold volume at day 29. (n=8 scaffolds/group)

### 3.7 Cell-assembled scaffolds containing ECFC alone had substantially remodelled into adipose tissue at day 29

Following μCT angiography, the scaffolds were excised and processed for H&E staining to assess scaffold remodeling and the distribution of perfused vessels within the implants (Figure 8). Interestingly, scaffolds containing ASCs, either alone or in combination with ECFCs, more frequently had perfused vessels (black due to the presence of the contrast agent) present in the central scaffold region, while the perfused vessels were primarily localized to the scaffold periphery in the ECFC alone group. Strikingly, while small pockets of adipocytes were observed in the periphery of the ASC+ECFC and ASC alone scaffolds, mature adipocytes were observed throughout the ECFC alone scaffolds, consistent with extensive remodeling into host-derived adipose tissue.

**Figure 8.**
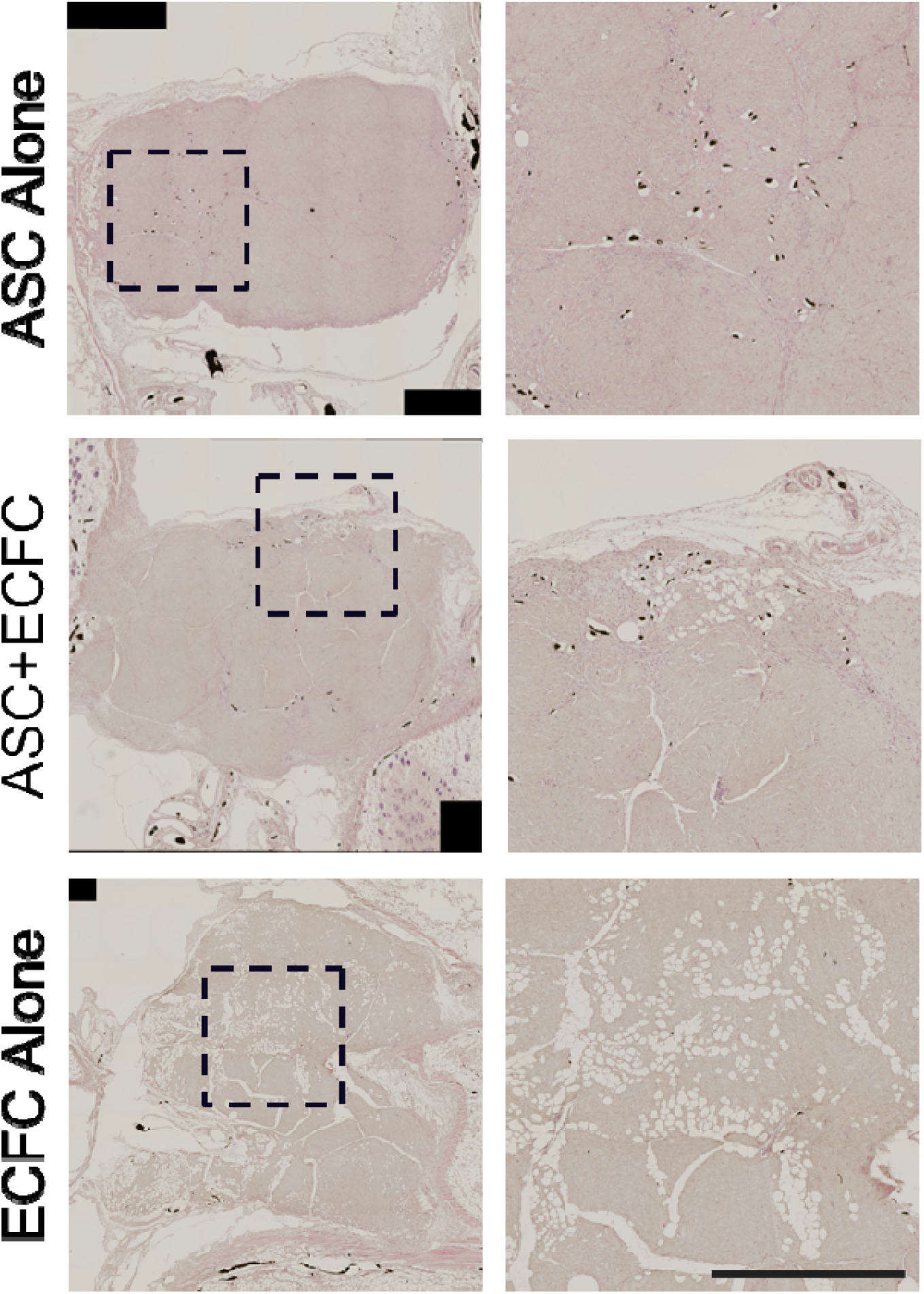
**Scaffolds containing ECFCs alone showed extensive remodeling, with adipocytes present in the central regions at day 29.** Representative images of the cell-assembled implants collected after μCT angiography at 29 days and stained with hematoxylin and eosin. Perfused vessels containing the contrast agent appear black, with more perfused vessels detected in the central regions of the scaffolds that contained ASCs based on qualitative analysis. Scale bars = 1 mm.

## 4 Discussion

DAT bioscaffolds have garnered great interest in the field of adipose tissue engineering given their demonstrated capacity to promote the adipogenic differentiation of ASCs *in vitro* and remodel into host-derived adipose tissue in pre-clinical *in vivo* models [9,14,45]. Although biomaterials-based approaches using the DAT alone as a volume filler are attractive from a translational perspective, fat formation can be enhanced by seeding the scaffolds with ASCs [14,46,47]. While originally targeted for their multipotent characteristics, it is now recognized that ASCs primarily function *in vivo* to promote host adipose tissue regeneration indirectly through paracrine-mediated mechanisms [48,49]. Angiogenesis and adipogenesis are closely interrelated, with an extensive network of capillaries required to support mature fat [50]. Importantly, ASCs produce a range of angiogenic factors [51], and can induce *in vivo* angiogenesis within adipose tissue via paracrine mechanisms [52].

Recognizing the potential advantages of bottom-up scaffold fabrication approaches, we recently developed a new method of generating “cell-assembled” scaffolds by applying ASC-seeded DAT microcarriers as building blocks to generate cohesive 3D engineered tissues that contain a high density of viable ASCs distributed throughout a dense adipose-derived ECM network [23] *In vivo* BLI showed that the localized retention of viable human ASCs was augmented at timepoints up to 7 days when the ASCs were implanted subcutaneously into *nu/nu* mice in the cell-assembled scaffolds compared to delivery on pre-fabricated DAT foams. While ASC retention was similar between the two platforms at 28 days, there was significantly enhanced host endothelial cell infiltration into the cell-assembled implants at day 28. Moreover, end-point μCT angiography revealed a striking reduction in vascular permeability in the cell-assembled scaffolds [23], suggesting that the cell-assembly approach more effectively harnessed the capacity of ASCs to stimulate functional vascular regeneration, which may be favorable for stable adipose tissue regeneration [21].

While the delivery of ASCs alone has shown promise for stimulating regeneration, there is evidence to support that combined cell therapies involving the co-delivery of cell populations with synergistic pro-regenerative capacities may be more effective [53]. Complementing ASCs, ECFCs represent a promising pro-angiogenic cell type that can form *de novo* vessels *in vivo* [54]. Additionally, ECFCs can modulate the recruitment of inflammatory cell populations involved in regeneration, as well as mural cells that can function to stabilize developing vessels [55].

Recognizing that our modular cell-assembly approach has the potential to generate 3D engineered tissues incorporating multiple cell populations, the primary objective of this study was to adapt our fabrication methods to enable the generation of scaffolds containing both human ASCs and ECFCs, and to characterize their properties relative to controls containing ASCs alone or ECFCs alone.

Preliminary testing revealed that the seeding efficiency could be better controlled by seeding the two cell types onto the DAT microcarriers within separate spinner flask bioreactors in their respective proliferation medium conditions. Notably, the ECFCs were seeded at twice the density of the ASCs to achieve a homogeneous distribution across the microcarrier surface at the end of the 24 h seeding period. While we were confident that we would be able to harness the capacity of the ASCs to secrete and remodel the ECM [56–58] to generate robust ASC+ECFC cell-assembled scaffolds, it was unclear whether we would be able to generate constructs containing ECFCs alone. Importantly, ECFCs can also secrete ECM components [59], as well as matrix metalloproteinases (MMPs) [60] that can contribute to remodelling of the DAT microcarriers.

While we were able to generate scaffolds containing ECFCs alone, there was macroscopically more variability in their level of cohesion, reflected in the varying levels of scaffold contraction in this group, and they were generally more fragile when being handled in comparison to the scaffolds containing ASCs. In the future, it would be interesting to compare the mechanical properties of the different scaffold groups on a more cellular level using sensitive techniques such as atomic force microscopy [61].

At the end of the 8-day assembly phase, all three scaffold groups were shown to contain cells distributed throughout their cross-section. While dense regions of ASCs were localized at the surface in the ASC+ECFC and ASC alone groups, the ECFCs were relatively evenly distributed throughout the scaffolds, with tubule-like structures observed in some regions in both the ASC+ECFC and ECFC alone groups. The PicoGreen and cell viability data supported that the ASCs were able to survive and proliferate within the scaffolds over time, consistent with their reported tolerance of ischemic conditions [62]. In contrast, flow cytometry suggested that cell viability was significantly lower in the ECFC alone scaffolds at day 8. This data could indicate that the ECFCs are less tolerant of the *in vitro* culture conditions required for scaffold assembly. However, a limitation of our approach is that ECFC viability may also have been negatively impacted by the 2-hour collagenase digestion step required to generate the single cell suspensions for flow analysis. Similarly, we were not able to obtain sufficient RNA yields for qPCR array analysis when we attempted to perform FACS to isolate the ECFC population from the scaffolds.

There is evidence that the co-culture of ASCs and ECFCs can be mutually beneficial for supporting cell survival and pro-angiogenic functionality [33,63]. For example, co-culturing ECFCs with ASCs has been reported to enhance tubule formation *in vitro*, with ASCs reported to have more potent effects compared to bone marrow- or endometrium-derived MSCs [34,64].

ASCs can act as pericytes, stabilizing the often transient or leaky vessels created by ECFCs [34]. In addition, co-culture of ASCs and ECFCs has been shown to be associated with increased levels of secreted pro-angiogenic factors including VEGF [63]. In the current study, the qPCR array analysis supports that co-culture with the ECFCs within the cell-assembled scaffolds modulated the ASC phenotype. Similar to our findings that showed a significant upregulation of ANGPT2 in the ASCs isolated from the ASC+ECFC scaffolds relative to the ASC alone group, Merfeld-Clauss *et al.* observed an upregulation in Ang-2 when human ASCs were co-cultured with endothelial cells [65]. In addition, Song *et al.* reported an increase in Ang-2 expression and decrease in TIMP1 expression when human tonsil-derived MSCs were cultured with ECFCs from cord blood [66].

The integration of *in vivo* cell tracking technologies into pre-clinical models is important for assessing the efficacy of cell delivery platforms at supporting the localized retention of viable cells, as well as understanding the underlying mechanisms of cell-based therapies [67]. In the current study, we successfully used a dual luciferase system to enable tracking of both the ASCs and ECFCs using *in vivo* BLI. A limitation is that the differing imaging timepoints makes it challenging to compare the kinetics of the two different cell populations. However, our strategy allowed us to quantitatively assess the effects of co-delivery on cell retention over time for both therapeutic cell populations. Interestingly, while ASC retention was similar in the ASC+ECFC and ASC alone scaffolds, co-delivery with ASCs was shown to significantly improve viable ECFC retention in the cell-assembled scaffolds at 4 weeks post-implantation. Our results are consistent with the findings of Song *et al.* who reported that co-delivery of human MSCs and ECFCs in combined cell spheroids improved ECFC retention, without impacting MSC retention, following intramuscular injection in a mouse hindlimb ischemia model based on *in vivo* fluorescence imaging [66]. Similarly, Kang *et al.* demonstrated using BLI that human ECFC retention could be enhanced by co-delivery in Matrigel with human bone marrow-derived mesenchymal progenitors in a murine hindlimb ischemia model [68]. The enhanced ECFC retention may be mediated through direct interactions between the co-delivered cell populations, which are important for enhancing ECFC tubule formation and stability *in vitro* [63], and/or via paracrine mechanisms [69].

Based on immunofluorescence staining for CD31 and μCT angiography, the vascular density was similar in the ASC+ECFC, ASC alone, and ECFC alone scaffolds at 29 days post-implantation. However, H&E staining of the scaffolds used for μCT angiography suggested that there may be some differences in the localization of the perfused vessels between the groups, with blinded analyses revealing that there were qualitatively more vessels containing Microfil in the central regions of ASC+ECFC and ASC alone scaffolds compared to those containing ECFCs alone. Notably, incorporation of any of the cell type(s) within the cell-assembled scaffolds was sufficient to stabilize the developing vasculature and prevent leaking of the contrast agent, which we previously observed in both ASC-seeded and unseeded pre-fabricated DAT foams [23].

Surprisingly, the H&E staining also revealed that large sections of the interior of the ECFC alone scaffolds had remodeled into adipose tissue, while adipocytes were only visualized along the periphery of the ASC+ECFC and ASC alone scaffolds. We had originally hypothesized that adipogenesis would be enhanced in the combined ASC+ECFC scaffolds, based on a previous study by Lin *et al.* that demonstrated enhanced human ASC engraftment and adipogenic differentiation when the ASCs were co-delivered subcutaneously in Matrigel with human ECFCs in immunodeficient mice [24]. One possibility is that the less cohesive nature of the ECFC alone scaffolds may have been favorable for the infiltration and/or differentiation of host adipogenic progenitor cells. To investigate this further, it would be interesting to assess the effects of shortening the 8-day cell-assembly phase for the ASC+ECFC and ASC alone groups to reduce the level of cohesion. In addition, extending the *in vivo* study to include longer timepoints would also provide valuable insight, with more extensive adipose tissue remodelling expected at timepoints ≥ 8 weeks [70,71].

## 5 Conclusions

In conclusion, this study successfully adapted our patented cell-assembly methods using a cell source other than ASCs for the first time, to enable the generation of scaffolds incorporating both human ASCs and ECFCs, as well as ECFCs alone, distributed throughout a cell-supportive DAT matrix. In this proof-of-concept study, we combined the ASC-seeded and ECFC-seeded microcarriers in a 1:1 ratio as a starting point for assessing the co-delivery of two cell types within the scaffolds. However, our modular fabrication strategy offers great flexibility. Varying ratios of the two cell types could easily be explored, and the methods could also be adapted to incorporate other cell types important for adipose tissue homeostasis, such as macrophages. It could also be interesting to explore the integration of bioprinting technologies to control the distribution of the ECFCs within the combined scaffolds as a strategy to enhance tubulogenesis *in vitro* and vascular regeneration *in vivo*. Overall, the cell-assembled bead foams represent a promising cell delivery platform that can be tuned to modulate regeneration. Co-delivery of ASCs and ECFCs was shown to influence the ASC phenotype *in vitro* and to enhance the localized retention of viable ECFCs *in vivo*. It would be interesting to explore the potential of the combined ASC+ECFC scaffolds in pre-clinical wound healing models in the future. In addition, future studies should probe the mechanisms through which the ECFC alone scaffolds promoted adipose tissue regeneration, including assessing whether the properties of the other scaffold groups could be tuned to induce a similar response.

## Data Availability Statement

The raw data supporting the conclusions of this manuscript will be made available by the authors, without undue reservation, to any qualified researcher.

## Acknowledgements

We thank our clinical collaborators, Dr. Aaron Grant, Dr. Damir Matic, and their teams, for their support with providing the adipose tissue samples used in this study. We also thank Dr. Kristin Chadwick and the London Regional Flow Cytometry Facility for technical guidance and support with the flow cytometry and cell sorting. In addition, Dr. Maria Drangova is acknowledged for providing access to the gravity perfusion system used for the angiography studies. Finally, our sincere thanks to Gillian Bell for her technical assistance and guidance. Funding for this study was provided by the Natural Sciences and Engineering Research Council of Canada (NSERC RGPIN-2017-0410) and the Canadian Institutes of Health Research (CIHR Institute of Musculoskeletal Health and Arthritis Priority Announcement #179858), with scholarship support for Sarah From provided through a Transdisciplinary Bone & Joint Training Award from the Bone and Joint Institute at Western University.

