## Supplementary Figure for "Co-delivery of human adipose-derived stromal cells and endothelial colony-forming cells in cell-assembled decellularized adipose tissue scaffolds for applications in soft tissue regeneration"

**Supplemental Figures**

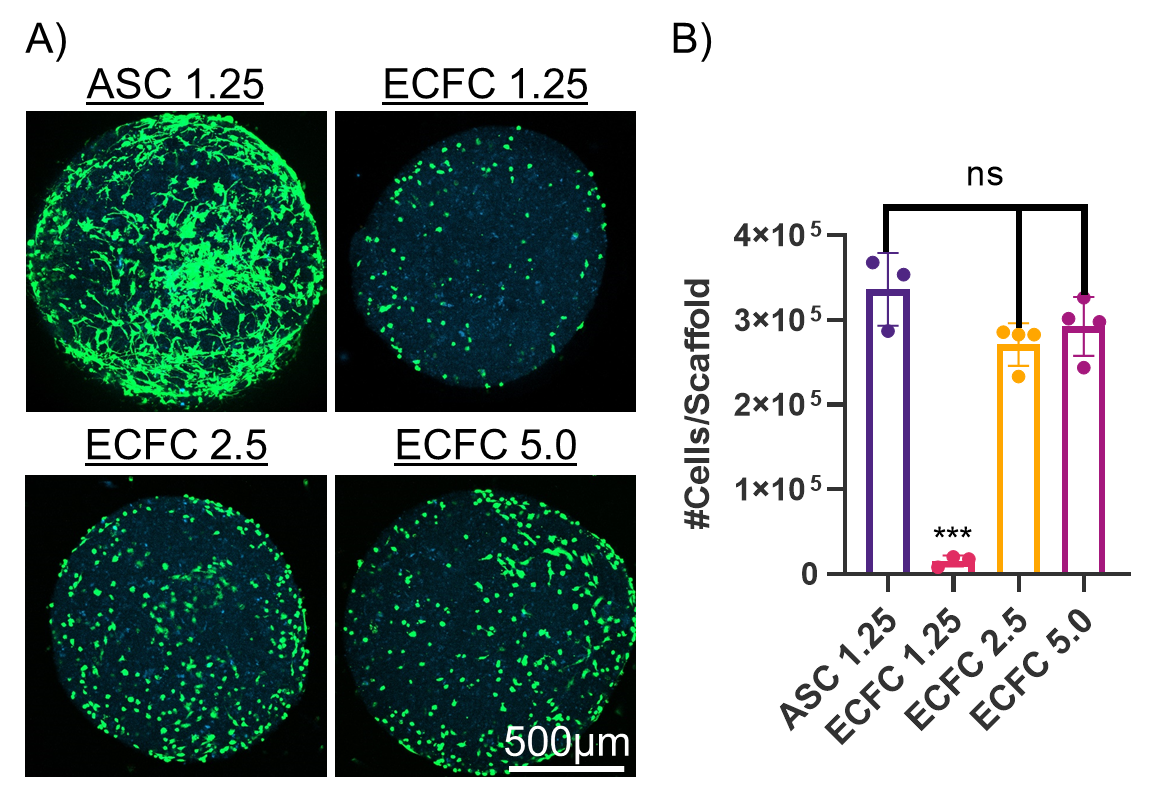

**Figure S1. ECFCs were seeded onto DAT microcarriers at a higher cell density (2.5 x 10^6^ cells/g) compared to the ASCs (1.25 x 10^6^ cells/g) to achieve a similar number of cells per scaffold.** (A) Representative Live/Dead® images at 24 h post-seeding. Live cells are shown in green (calcein AM) and blue is ECM autofluorescence. (B) At 24 h post-seeding, the dsDNA content of moulded scaffolds fabricated with the cell-seeded microcarriers was measured via the PicoGreen™ assay and converted to cell number using a standard curve (n=3, N=3-4, One-way ANOVA). ***p<0.005.

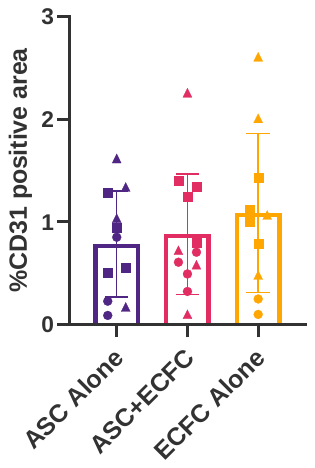

Figure S2. No significant differences were observed in the percentage of CD31 positive pixels per scaffold area between the scaffold groups at day 29 post-implantation. Images were taken of 4-6 fields of view/slide and 2-3 slides/scaffold (n=7-8 scaffolds/group).

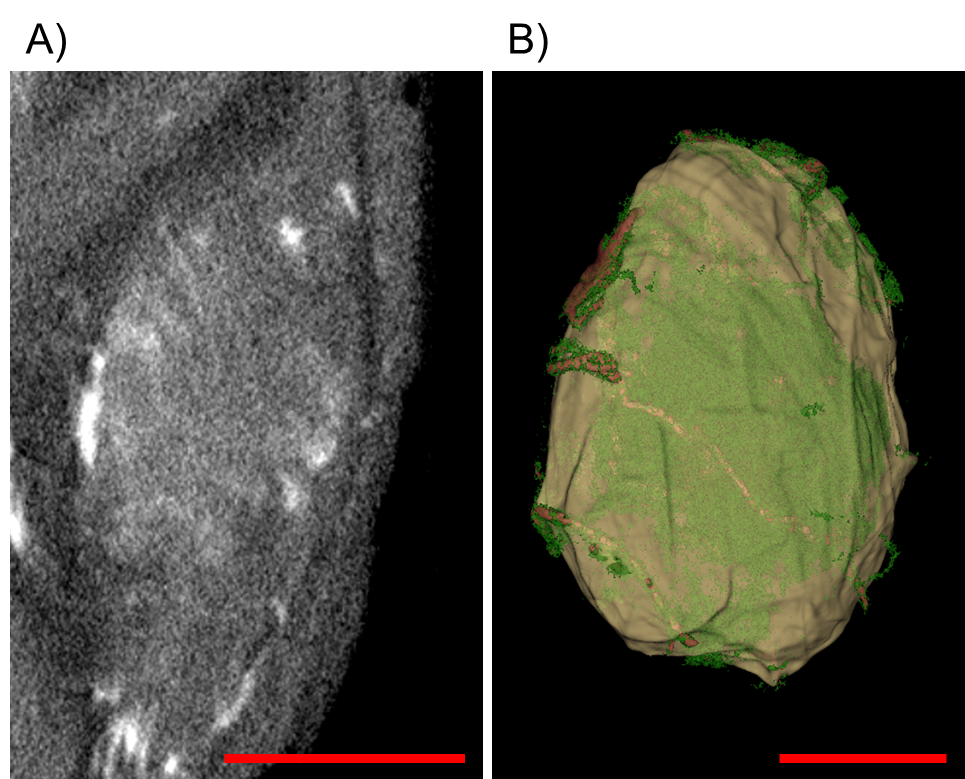

Figure S3. Unseeded scaffolds consistently showed leaking of the contrast agent, which was not detected in any other group. Representative images of perfused unseeded scaffold after 29 days *in vivo.* (A) Maximum intensity projections of 300 µm section from the middle of the implanted scaffold show leaking of the contrast agent. (B) 3-D model of the scaffolds (shown in beige), with perfused vessels shown in red and the leaked contrast agent detected within the scaffold shown in green. Scale bars = 2.5 mm.
